# Eco-evolutionary feedbacks generate bistability in population persistence under gradual environmental change

**DOI:** 10.64898/2026.08.05.743160

**Authors:** Kuangyi Xu, Hao Shen

## Abstract

Understanding how populations persist in gradually deteriorating environments through evolution is a central question in ecology and evolutionary biology. Previous studies have primarily focused on identifying the critical rate of environmental change beyond which extinction is certain. However, the existence of a viable equilibrium when the rate is below the threshold does not guarantee that a population can survive the transient dynamics to reach it. Using a quantitative genetic model that explicitly incorporates feedback among population size, genetic variance, and mean trait evolution, we show that population persistence can exhibit bistability when the rate of environmental change is below the extinction threshold. Specifically, extinction still occurs if the initial population size and genetic variance fall below a critical level. The initial state also influences the eco-evolutionary dynamics, such that a temporary increase or decline in population size and/or genetic variance does not necessarily predict the ultimate fate of the population. Therefore, in addition to estimating the critical rate of environmental change for extinction, characterizing current population size, genetic variation, and the degree of maladaptation may improve predictions of extinction risk in deteriorating environments.

## 1 Introduction

Gradual environmental change is increasingly threatening the persistence of natural populations, with many species exhibiting demographic declines or local extinctions in response to ongoing climate change (Parmesan, 2006; Urban, 2015). Extinction may be avoided if populations can evolve rapidly enough to maintain non-negative population growth despite continual environmental deterioration (Alexander et al., 2014; Bell, 2017; Gonzalez, Ronce, et al., 2013; Hoffmann and Sgró, 2011). Consequently, understanding the conditions under which evolution enables long-term population persistence in changing environments has attracted considerable theoretical interest (reviewed in Klausmeier et al., 2020; Kopp and Matuszewski, 2014; Uecker et al., 2026) and has important implications for biodiversity conservation and the management of drug resistance (Alexander et al., 2014; Radchuk et al., 2019).

Existing theory has largely focused on identifying the condition when population can persist by tracking a continuously moving phenotypic optimum while maintaining non-negative population growth (Bürger and Lynch, 1995; Klausmeier et al., 2020; Lande and Shannon, 1996; Lynch and Lande, 1993; Matuszewski et al., 2015; Osmond and Klausmeier, 2017). These studies predict the existence of a critical rate of environmental change, above which populations inevitably go extinct. Extinction occurs either because mean population growth becomes negative although the population mean trait value can trail the moving optimum by a constant lag (demographic constraint; Gomulkiewicz and Houle, 2009), or because evolution cannot keep pace with environmental change, causing the lag to increase without bound (an evolutionary constraint; Osmond and Klausmeier, 2017). Correspondingly, estimates of critical environmental change rates have been proposed as a means of assessing extinction risk under ongoing climate change (Gienapp et al., 2013; Willi and Hoffmann, 2009).

Nevertheless, the existence of a viable equilibrium does not guarantee that a population can persist regardless of its initial conditions. As pointed out qualitatively by Bürger and Lynch (1995), an excessively large phenotypic lag can cause population decline, leading to stronger genetic drift and diminished genetic variance. The resulting loss of evolutionary potential further slows adaptation and intensifies demographic decline, potentially driving the population toward extinction. Consequently, extinction may also represent a locally stable equilibrium. This means that populations facing identical environmental conditions may experience divergent ultimate fates depending on their initial states.

This distinction between equilibrium existence and its accessibility has received relatively little attention. Although models of evolutionary rescue following abrupt environmental shifts have examined how nonequilibrium eco-evolutionary dynamics can determine extinction versus rescue (Nordstrom et al., 2023; Xu et al., 2023), models considering persistence under gradual environmental changes have rarely done so. Klausmeier et al., 2020 illustrated that when selective death depends on population density, bistability can occur and population persistence will depend on the initial density when the rate of environmental change is intermediate. This example establishes that equilibrium accessibility can matter, but it remains unclear whether bistability can arise from the canonical eco-evolutionary feedback among demographic decline, genetic drift, genetic variance, and adaptation, and how a population’s ultimate fate depends on its initial eco-evolutionary state.

Here, we develop a logistic-growth model that explicitly incorporates feedback among mean trait evolution, genetic variance, and population size under a gradually changing environment. We show that while a critical rate of environmental change remains a central boundary separating sustainable adaptation from inevitable extinction, persistence below this critical rate is far from guaranteed. Instead, when populations initially lag behind the environmental optimum, the system often exhibits bistability: populations with insufficient initial population size or genetic variance ultimately go extinct, even when it exhibits temporary increases in both quantities. We further demonstrate the biological importance of this dependence on initial conditions by examining several relevant scenarios.

Our results suggest that more reliable predictions of population persistence or extinction may require information not only about the rate of environmental change but also about the population’s initial demographic and evolutionary state.

## 2 Model

We consider a haploid population with overlapping generations, with either asexual reproduction or random mating, subject to density-dependent population growth regulation. Adaptation occurs through the evolution of a quantitative trait, *z*, which follows a Gaussian distribution with mean trait value *µ*(*t*) and variance *V* (*t*) at time *t*. For simplicity, we assume that the trait is strictly heritable and polygenic, governed by numerous loci of equal minor additive effect. Individuals experience Gaussian stabilizing selection, and the fitness function in continuous time is *g*(*z*) = *r* − (*z* − *θ*(*t*))^2^*/*2*ω*^2^, where *r* is the maximum growth rate, and *ω*^2^ determines the strength of stabilizing selection around the phenotypic optimum *θ*(*t*). We assume the selection is weak, such that the width of the fitness landscape is large compared to the phenotypic variance (*ω*^2^ ≫ *V* (*t*)).

We define the lag between the population mean and the phenotypic optimum as Δ(*t*) = *θ*(*t*) − *µ*(*t*) and let the population size be *N* (*t*). The population is assumed to be under logistic growth with mean intrinsic growth rate given by

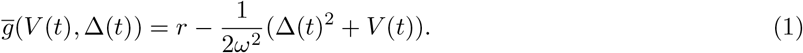

The two terms in the parenthesis represent the fitness reduction due to the mismatch between the mean trait value and the optimum (lag load) and phenotypic variation (variance load), respectively. The change of population size *N* (*t*) is

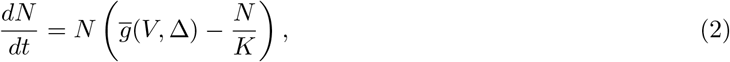

where *K* determines the strength of density regulation.

We are interested in whether the population can persist through adaptive tracking. Extending previous models (Bürger and Lynch, 1995), we assume that the mean trait value is initially 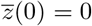 without loss of generality, and that the phenotypic optimum changes linearly over time as *θ*(*t*) = *θ*_0_ + *vt*. This formulation incorporates both an initial abrupt environmental shift of magnitude *θ*_0_ and a subsequent gradual movement of the phenotypic optimum at a constant rate *v*.

We characterize the system by tracking the dynamics of Δ(*t*), *V* (*t*) and *N* (*t*). For notational simplicity, we omit the explicit time argument (*t*) for all dynamic variables hereafter.

The rate of evolution in the mean trait value is (Lande, 1979)

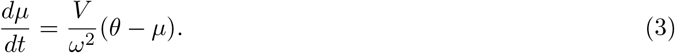

Since *θ*(*t*) = *θ*_0_ + *vt*, the dynamics of the lag are given by

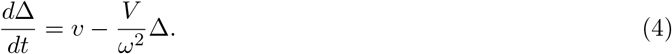

The change of genetic variance is (Kimura, 1965b; Lande, 1993)

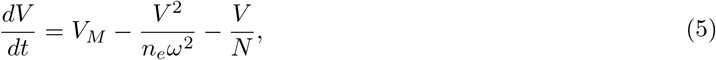

where *V*_*M*_ is the input of genetic variance via mutation, and the second and third terms represent the reduction of genetic variance by selection and genetic drift, respectively. *n*_*e*_ is the effective number of recombining factors, and when loci underlying trait *z* are freely recombining, *n*_*e*_ is equal to the number of loci.Particularly, for an asexually reproducing population, *n*_*e*_ = 1. It is important to note that, for a sexually reproducing population, Equation (5) is an approximation under several assumptions. First, it assumes that dynamics of linkage disequilibrium (LD) are much faster than allele frequency dynamics, such that LD structure among recombining factors is always in quasi-linkage equilibrium (QLE) (Kimura, 1965a). Second, Equation (5) ignores the contribution of LD under QLE to the additive genetic variance, which is a natural consequence of the weak selection assumption. Due to the interplay of selection and recombination, the rigorous expression for the genetic variance dynamics can be more complicated than Equation (5) (Bulmer, 1971; Lande, 1975; Negm and Veller, 2026; Turelli and Barton, 1990), and for a more detailed analysis of the influence of LD, we refer the readers to Negm and Veller, 2026.

We denote the phenotypic lag, genetic variance and population size at the onset of environmental change (*t* = 0) by *V*_0_, *N*_0_ and Δ_0_, respectively. Note that Δ_0_ may be nonzero if the environmental change leads to an abrupt shift in the phenotypic optimum, or because the mean trait value does not match the original phenotypic optimum prior to the environmental change, for example, as a result of genetic drift or ongoing gradual environmental change.

## 3 Results

### 3.1 Critical rate and viable equilibrium

In Appendix S1, we prove that there exists a critical moving rate, *v*_*c*_, above which population extinction is certain for any initial states, characterized by

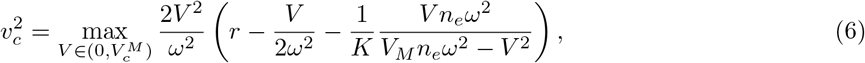

where 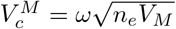 is the equilibrium genetic variance in infinitely large population. The critical moving rate increases with a larger mutation input of genetic variance (boundary curve in Figure 1), while it tends to peak at intermediate strength of selection (Figure 2a), qualitatively consistent with previous results (Xu and Osmond, 2026). The selection strength maximizing *v*_*c*_ is lower when the mutation input *V*_*M*_ is larger and density regulation (1*/K*) is weaker (compare the blue line with other lines in Figure 2a).

**Figure 1:**
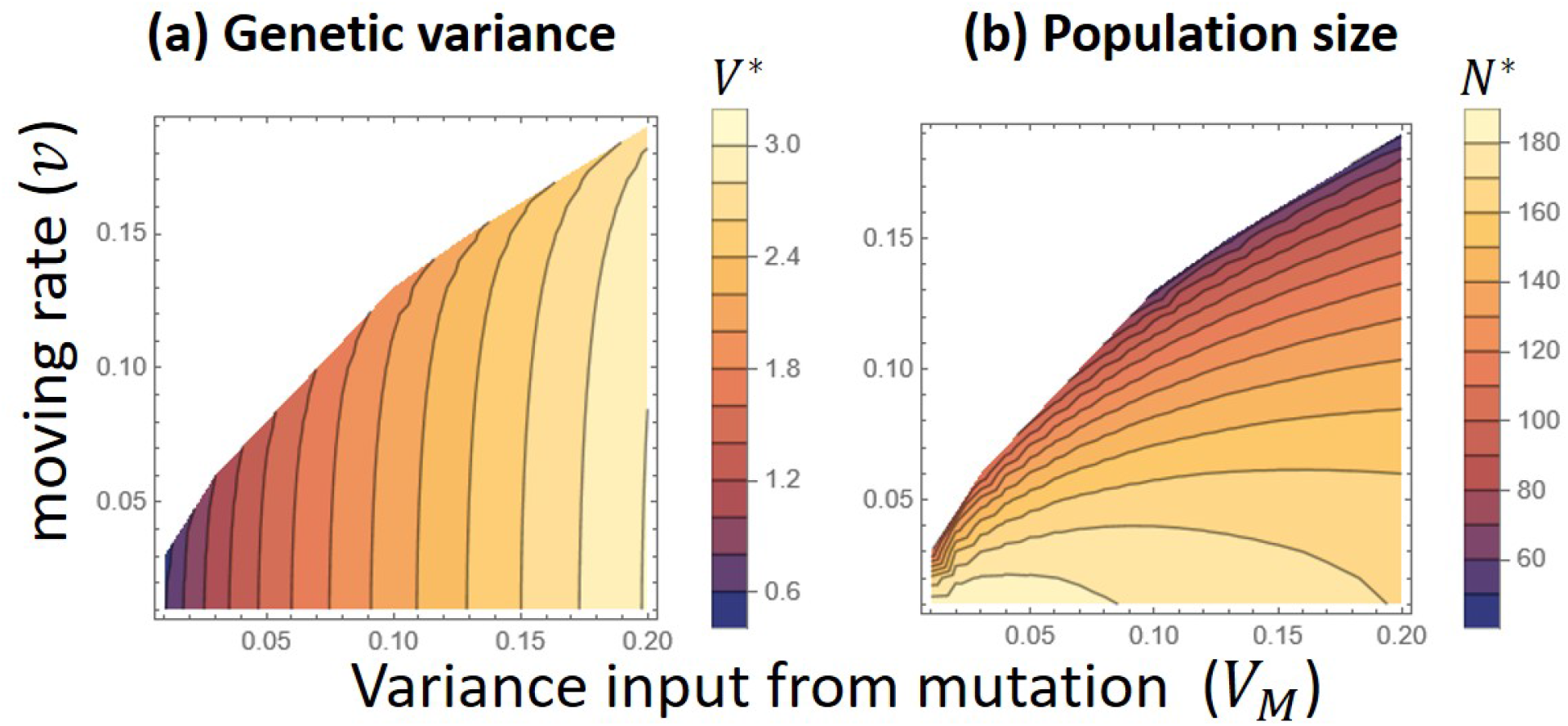
Impacts of the mutation input (*V*_*M*_ ) on the critical moving rate,*v*_*c*_, above which extinction is certain (boundary of colored area), as well as the equilibrium genetic variance and population size when the moving rate *v* is below *v*_*c*_. Results are obtained based on Equations (6)-(8). Parameters: *r* = 0.2, *ω*^2^ = 50, *V*_*M*_ = 0.1, *K* = 1000, *n*_*e*_ = 1.

**Figure 2:**
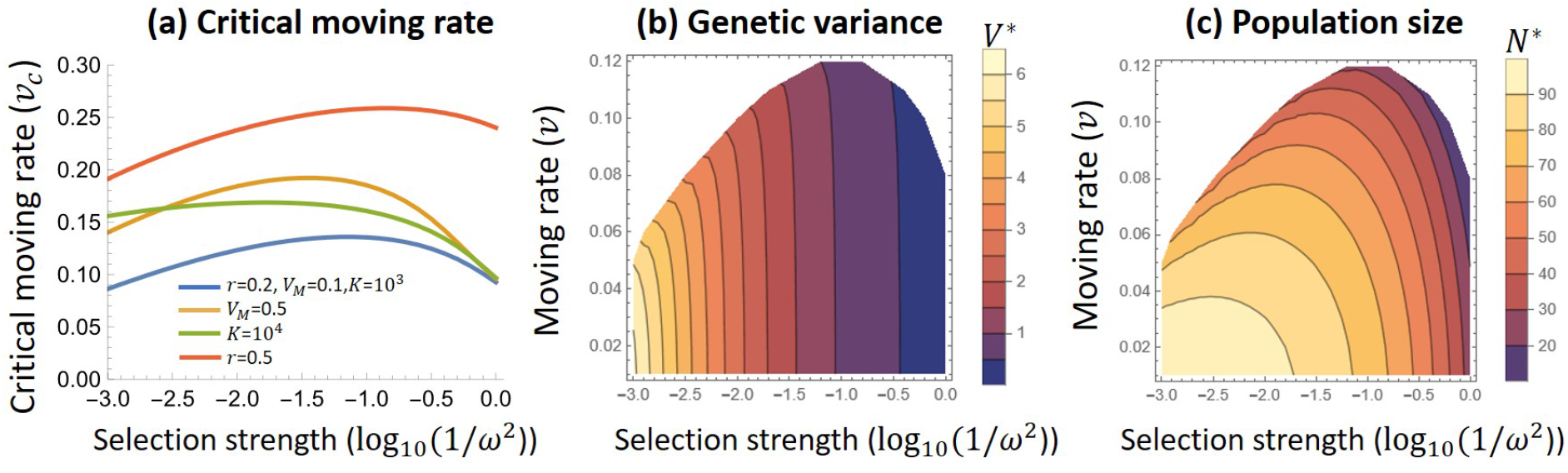
**(a):** Impacts of strength of stabilizing selection (1*/ω*^2^) on the critical moving rate (*v*_*c*_) above which extinction is certain. Blue line is taken as the standard parameter and other lines differ from it in one of the parameters. **(b) &( c)**: Impacts of selection strength and the moving rate on the equilibrium genetic variance and population size (*v*_*c*_ is the boundary of the colored region). Results are obtained based on Equations (6)-(8). Parameters: *r* = 0.2, *V*_*M*_ = 0.1, *K* = 1000, *n*_*e*_ = 1.

When the moving rate is below the critical rate (*v* < *v*_*c*_), there exist two equilibrium states: 1) extinction (*N* = 0), and 2) a viable equilibrium at which populations can persist by tracking the moving optimum with a constant lag. Both equilibria are shown to be locally stable (see Appendix S2 for the proof). Thus, whether the population can persist or not depends on its initial eco-evolutionary state.

At the viable equilibrium, the lag and the population size are

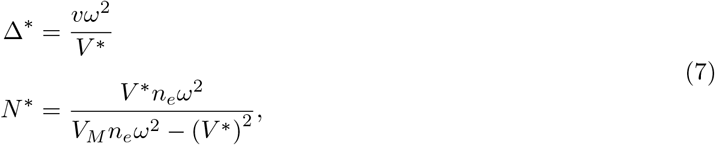

where the equilibrium genetic variance *V* ^∗^ is a solution to the equation

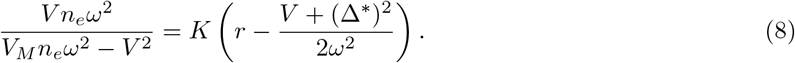

The left-hand side is the population size required to maintain a specific phenotypic variance, and the right-hand side is the population size constrained by fitness shaped by moving phenotypic optimum and density dependence.

The equilibrium genetic variance slightly decreases with faster moving rate, but is primarily determined by the magnitude of mutation input *V*_*M*_ and selection strength 1*/ω*^2^ (Figures 1a and 2b). In contrast, the equilibrium population size often peaks at intermediate levels of mutation input (and thus intermediate genetic variance) (Figure 1b) and at intermediate strengths of stabilizing selection (Figure 2c). This pattern arises because low genetic variance or weak stabilizing selection slows adaptation, resulting in a large lag load, whereas high genetic variance or strong stabilizing selection generates a large variance load.

### 3.2 Impacts of initial eco-evolutionary states

Even when the rate of environmental change is below the critical rate (*v* < *v*_*c*_), if *v* is relatively high and the population initially exhibits a lag (Δ_0_ > 0), the population may still go extinct if its initial genetic variance and population size fall below a critical threshold (the region below the blue curve in Figure 3). The minimum initial genetic variance required for persistence decreases as the initial population size increases. The region of the initial-condition space leading to extinction expands rapidly as either the initial lag or the rate of environmental change increases (Figure 3).

**Figure 3:**
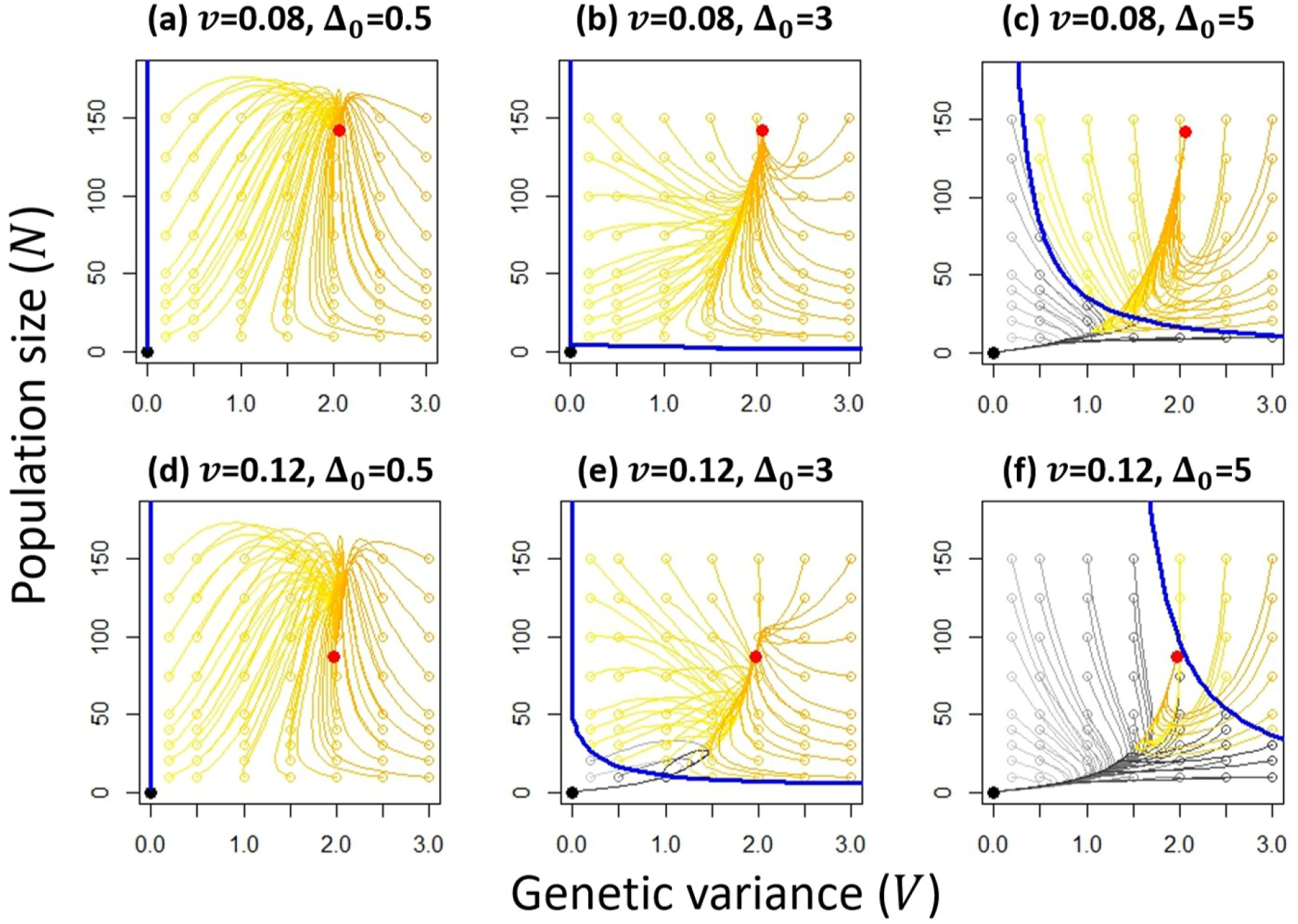
Impacts of initial conditions on population final survival status, as well as eco-evolutionary trajectories of population size and genetic variance. Open circles indicate the starting state of each trajectory. The blue curve marks the basin boundary of initial population size (*N*_0_) and initial genetic variance (*V*_0_). Populations initialized below this boundary will go extinct (gray trajectories). Populations starting above the boundary persist (yellow-to-orange trajectories) and eventually evolve toward a stable steady state (red dot). Note that the trajectories of persistent populations may cross the boundary since the lag also changes (not shown). The three columns show results as the initial lag Δ_0_ increases from low to high. Results are obtained based on Equations (2)-(5). Parameters: *r* = 0.2, *ω*^2^ = 50, *V*_*M*_ = 0.1, *K* = 1000, *n*_*e*_ = 1.

The initial eco-evolutionary state of a population also influences the dynamics towards the equilibrium. When the initial lag Δ_0_ is much smaller than the equilibrium lag, since the lag load gradually increases over time, the population size often exhibits an initial increase followed by a decline on the way to equilibrium across a broad range of initial population sizes and initial genetic variances (Figures 3a, d). Conversely, when Δ_0_ is much larger than the equilibrium lag, the population may often undergo a temporary decline in both demography and genetic variance, followed by a subsequent rebound (orange trajectories in Figures 3c, f). When the initial lag is close to the equilibrium lag, interestingly, a population may ultimately go extinct even when it exhibits a temporary increase in both population size and genetic variance (e.g., gray trajectories in Figure 3e).

### 3.3 Example scenarios

In the above section, the initial genetic variance, population size, and lag are varied as independent parameters for generality. In reality, however, these variables may be interdependent and jointly determined by biological and environmental processes. Here, we examine how bistability influences population persistence under three biologically relevant scenarios.

One biologically realistic scenario is that the initial genetic variance, *V*_0_, is at mutation–selection–drift balance and therefore depends on the initial population size *N*_0_. In this case, persistence is possible only when the initial population size is large enough, which maintains a sufficient large genetic variance. Although stronger selection increases the rate of evolution, the minimum population size required for persistence increases with the strength of stabilizing selection, and persistence is impossible when selection is too strong (Figure 4a). In other words, populations subject to stronger stabilizing selection are less likely to persist under environmental change. This is because selection reduces genetic variance, particularly when population sizes are small. Additionally, the minimum initial population size required for persistence also increases with greater initial lag (Δ_0_) and the rate of environmental change (*v*) (Figure 4a).

**Figure 4:**
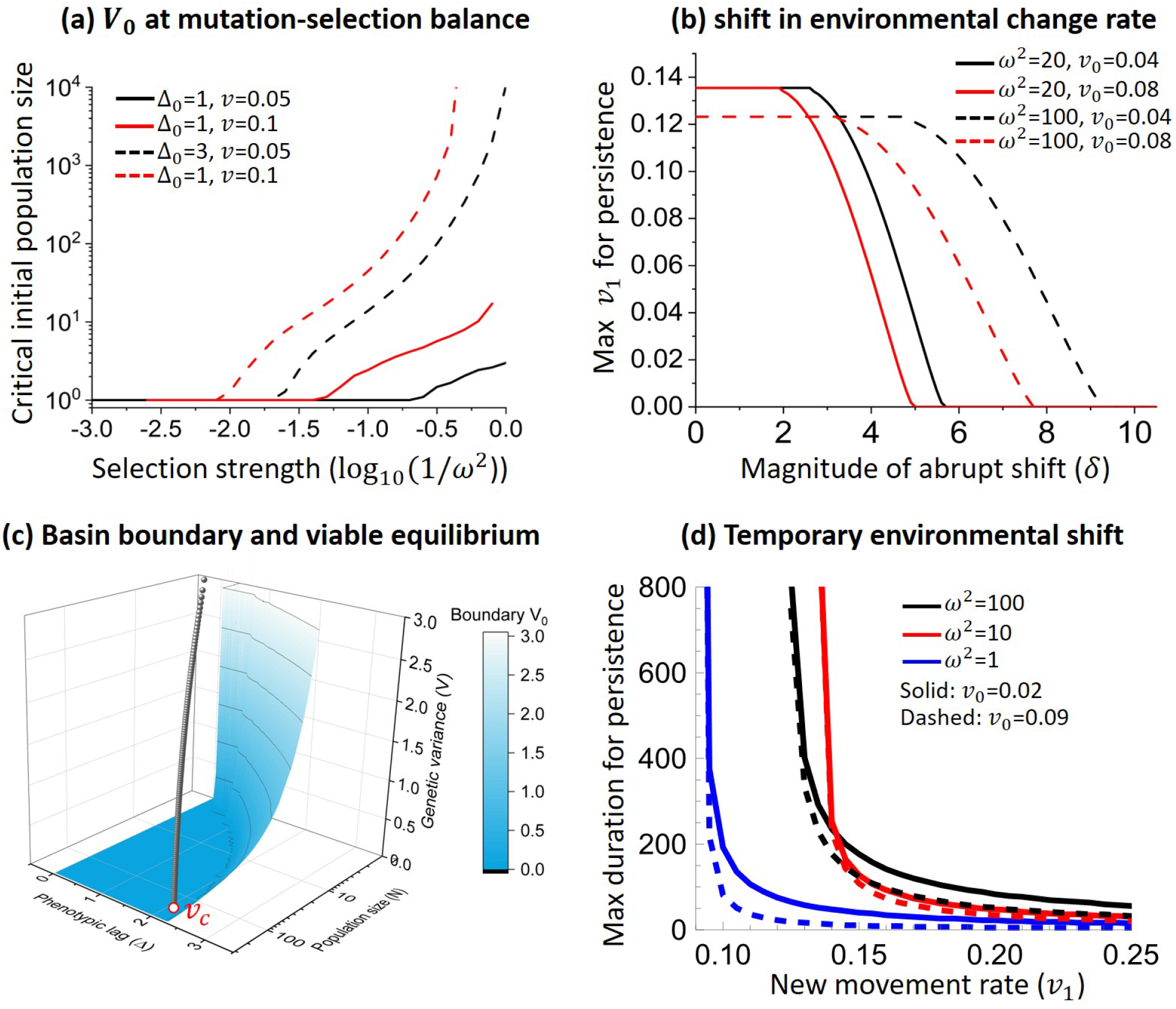
Impacts of bistability on population persistence under three biologically relevant scenarios. **(a)**: When initial genetic variance *V*_0_ is under mutation-selection-drift balance. There is a minimum initial population size *N*_0_ for populations to persist in a gradually changing environment. The solid curves are truncated since *v*_*c*_ depends on selection strength and *v* > *v*_*c*_ when selection strength exceeds a certain level. The dashed curves go to infinity, meaning that persistence is no longer possible when selection is too strong even when *v* < *v*_*c*_. **(b)**: A population initially at the viable equilibrium under moving rate *v*_0_ experiences an environmental change, which alters the moving rate from *v*_1_ and/or causes an abrupt shift in phenotypic optimum *δ*. The curve shows the maximum *v*_1_ allowing persistence, depending on the change of phenotypic lag. **(c)**: The blue plane marks the basin boundary for which a population starting above it can persist as long as *v* < *v*_*c*_. The red curve shows the equilibrium states when the moving rate *v* changes from 0 to *v*_*c*_, which will hit the boundary as *v* → *v*_*c*_. **(d)**: A population initially at equilibrium under an environmental change rate *v*_0_ experiences a temporary environmental shift that changes the rate to *v*_1_ for a duration *T* . The figure shows the maximum duration that still allows the population to persist after the rate returns to *v*_0_. Note that the maximum duration will go to infinity as *v*_1_ approaches *v*_*c*_, and *v*_*c*_ changes with selection strength *ω*^2^. For all panels, unless otherwise specified, parameters are *r* = 0.2, *ω*^2^ = 50, *V*_*M*_ = 0.1, *K* = 1000, *n*_*e*_ = 1.

A second biologically relevant scenario is that a population initially persists under a gradually changing environment but subsequently experiences further environmental change. Accordingly, we consider a population that is initially at equilibrium under an optimum moving rate *v*_0_. An environmental shift then changes the moving rate from *v*_0_ to *v*_1_, and may also generate an abrupt shift in the phenotypic optimum of magnitude *δ*. The population can persist only when both *v*_1_ and *δ* remain below threshold values. In general, persistence is more likely when the original moving rate, *v*_0_, is lower (compare the red and black lines in Figure 4b) and when selection is weaker (compare the solid and dashed lines in Figure 4b).

The maximum rate *v*_1_ that still allows persistence is lower as the abrupt shift *δ* becomes larger, and when *δ* is too large, persistence becomes impossible for any values of *v*_1_, and *vice versa* (Figure 4b). In particular, when there is no abrupt shift (i.e., *δ* = 0), populations that are initially at equilibrium under any original moving rate *v*_0_ can always persist under the new rate *v*_1_, as long as *v*_1_ is below the extinction threshold *v*_*c*_. This is because the equilibrium state under *v*_0_ will always lie above the basin boundary of persistence under *v*_1_ < *v*_*c*_ (Figure 4c), so that a population starting at equilibrium state under *v*_0_ can always reach the equilibrium state under *v*_1_.

It is possible that an environmental shift is temporary. Therefore, the third scenario consider an equilibrium population initially under a moving rate *v*_0_. The rate increases from *v*_0_ to *v*_1_ for a duration *T* and then returns to *v*_0_. Even when *v*_1_ exceeds the extinction threshold rate *v*_*c*_, populations may still persist if the duration *T* is sufficiently short. However, when *T* is too long, bistability predicts that populations may go extinct even after environmental conditions recover, a phenomenon known as “extinction debt” (Osmond and Klausmeier, 2017). The maximum duration that still allows persistence decreases rapidly as *v*_1_ increases from the threshold rate *v*_*c*_ (Figure 4d). Although stronger selection allows faster trait evolution under the new rate, it may often reduce the maximum duration due to increase of variance load, unless the new rate is slightly above the critical threshold (compare lines with *ω*^2^ = 100, 10, 1 in Figure 4d).

## 4 Discussion

Using a deterministic eco-evolutionary model, this study investigates how feedbacks among demography, trait value, and genetic variance influence population persistence under gradual environmental change through the evolution of a quantitative trait. In contrast to previous theoretical results where persistence is guaranteed provided the rate of environmental change is below a critical threshold, incorporation of eco-evolutionary feedback shows that persistence additionally requires sufficiently large initial population size and genetic variance, along with a sufficiently small initial lag. The initial state also influences the transient dynamics towards the viable equilibrium, such that temporary changes in population size or genetic variance may not predict their ultimate fate.

Our finding that population persistence depends on the initial state is analogous to the tipping point found in Osmond and Klausmeier, 2017 and the bistability example in Klausmeier et al., 2020. However, the underlying mechanisms slightly differ. The tipping point arises because under a non-quadratic fitness function, the rate of evolution declines once the lag exceeds a critical threshold, while in Klausmeier et al., 2020, selection is weaker in smaller populations due to density-dependent death rates. As a result, populations with too large lag or too low density will go extinct due to slow evolution. In our model, too large lag causes reduction of population size and thus in genetic variance, leading to slower evolution in future generations (i.e., extinction vortex).

Due to the impact of the initial state, population persistence depends on historical environmental conditions. For a population already persisting under gradual environmental change, we find that it can continue to persist after the rate of environmental change suddenly increases to a higher level, provided that the new rate remains below the extinction threshold. However, an abrupt shift in the phenotypic optimum accompanying the rate change can greatly reduce the maximum new rate that still allows persistence, especially when selection is strong (Figure 4b). Moreover, a temporary rise in the rate of environmental change exceeding the extinction threshold can drive the population to extinction, even when the rate returns to the original value, similar to the result in Osmond and Klausmeier, 2017. The maximum duration of this temporary increase that still allows persistence after recovery declines rapidly as the new rate increases (Figure 4d).

### 4.1 Implication for empirical studies

Previous models of evolutionary rescue have typically assumed that populations are at their ecological and evolutionary equilibrium before environmental change occurs. In reality, however, populations are unlikely to be exactly at equilibrium even in a constant environment, as demographic and evolutionary stochasticity continuously generates perturbations in population size, mean trait value, and genetic variance (Lande, 1993; Xu et al., 2023). Moreover, natural environments themselves often fluctuate, further displacing populations from equilibrium. Consequently, even replicate populations under the same stable environment may vary in their initial states, which can critically determine their ultimate fate (Klausmeier et al., 2020; Xu et al., 2023).

Recording demographic dynamics has been an important indicator of whether evolutionary rescue has occurred in nature (Anstett et al., 2026). We find that demographic trajectory during evolutionary rescue may depend on the initial conditions. While some rescued populations exhibit the typical U-shaped trajectory, others may instead experience a temporary increase followed by a decline in population size, even when the initial population size exceeds the equilibrium value (Figure 3a,d). Importantly, a transient increase or decrease in population size or genetic variance does not necessarily predict a population’s ultimate fate. Depending on the initial eco-evolutionary state, populations that initially appear to recover in demography and genetic variation may ultimately go extinct, whereas those showing temporary decline may eventually persist (Figure 3). Therefore, estimates of the current demographic and evolutionary potential could improve assessments of extinction risk under environmental change.

Previous experiments have demonstrated that faster rates of environmental change and stochastic fluctuations in those rates reduce the probability of evolutionary rescue in gradually deteriorating environments (Hao et al., 2015; Lindsey et al., 2013; Zhou and Zhang, 2024). However, in these experiments, replicate populations were typically established under similar initial conditions. Rescue experiments imposing an abrupt environmental shift have confirmed the theoretical prediction that evolutionary rescue is promoted by larger initial population sizes (Bell and Gonzalez, 2009; Ramsayer et al., 2013; Samani and Bell, 2010) and greater standing genetic variation (Agashe et al., 2011; Lachapelle and Bell, 2012). Extending these experiments to gradually deteriorating environments while varying initial demographic and initial genetic conditions would directly test our prediction that population persistence depends critically on initial conditions.

Our model predicts that populations undergoing faster environmental deterioration are less likely to survive a subsequent abrupt environmental shift (Figure 4b). However, experimental studies have found that prior exposure to mild stress or a gradually deteriorating environment can facilitate rescue following an abrupt exposure to high levels of stress (Bell and Gonzalez, 2011; Gonzalez and Bell, 2013; Samani and Bell, 2010). This inconsistency is likely due to differences in the genetic architecture of adaptation. Our model assumes adaptation from many loci of small effect, whereas the experimental systems involve resistance evolution at a few major-effect loci. In the latter case, prior exposure to mild stress increases the likelihood that high-resistance mutations arise from intermediate-resistance genotypes (Osmond et al., 2020). In fact, prior stress exposure can increase the extinction likelihood when challenged with a qualitatively different stress (Lachapelle et al., 2017; Samani and Bell, 2016).

### 4.2 Limitations and future directions

Following previous models (e.g., Bell, 2017; Lande and Shannon, 1996), we adopt a deterministic model, which ignores stochasticity in demographic and evolutionary changes. Incorporating stochasticity in population growth and the evolution of mean trait value and genetic variance should not qualitatively change the results, although the critical moving rate will be lower than the deterministic model predicts (Bürger and Lynch, 1995; Lynch and Lande, 1993), and the basin parameter space of initial conditions towards extinction will be larger.

Also, as many previous models did, we assume a Gaussian fitness function, such that the rate of evolution is proportional to the lag (Equation 3), while the reduction in genetic variance caused by selection is independent of the lag (the second term in Equation 5). Under other fitness functions, however, the rate of evolution may exhibit non-linear dependency (Klausmeier et al., 2020; Osmond and Klausmeier, 2017; Xu and Osmond, 2026), and the reduction in genetic variance caused by selection may depend on the lag, such that a larger lag may lead to slower evolution and faster depletion of genetic variance by selection. Moreover, the rate of environmental change may also influence genetic variance by altering the genetic basis of polygenic adaptation (Höllinger et al., 2019; Kopp and Matuszewski, 2014; Kopp and Hermisson, 2009; Matuszewski et al., 2015; Pahujani et al., 2026). Therefore, our model may be better viewed as a proof-of-concept model (Servedio et al., 2014). Although we expect our main conclusions to remain qualitatively robust, incorporating more general fitness functions and more detailed genetic basis underlying adaptation may reveal richer forms of eco-evolutionary feedback and dynamics.

## Acknowledgments

We would like to thank Matthew Osmond, Walid Mawass and members of the John Novembre lab for their
feedback.

## Supplementary materials

### S1 Equilibrium and critical moving rate

We write the full coupled dynamical system as

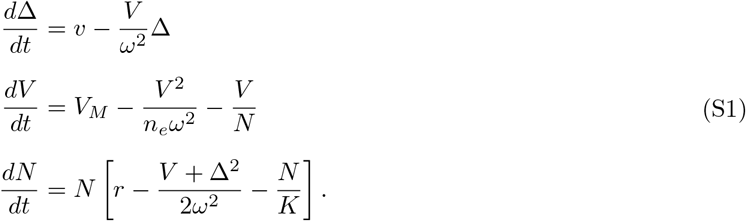

Notice that in all cases, 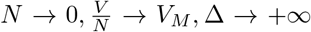 is a stable equilibrium, thus an extinction vortex always exists if the initial Δ is too large and *N* and *V* are too small. The question is when the initial parameter is proper, can an evolutionary rescue happen. To be more specific, we would like to know when would a positive stable equilibrium (Δ^∗^, *V* ^∗^, *N* ^∗^) exists.

To solve for the equilibrium, by setting 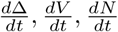 to be zero, we get

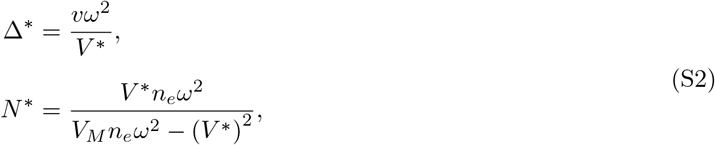

where the *V* ^∗^ is a solution to the equation

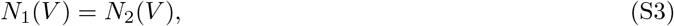

where 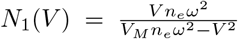 is the population size required to maintain a specific phenotypic variance, and 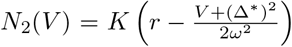 is the population size constrained by fitness shaped by moving phenotypic optimum and density dependence. At the equilibrium, we have the following equation

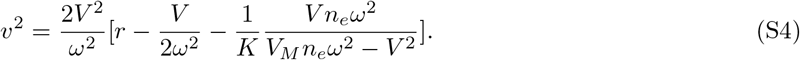

We now look at the range of *V* . Obviously, we would require *v*^2^ ≥ 0 and *V*_*M*_ *n*_*e*_*ω*^2^ − *V* ^2^ ≥ 0, the second inequality gives 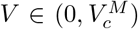 where 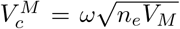. Notice that for 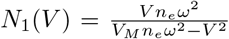 monotonically increases with *V*, the function 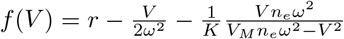 thus monotonically decreases from *r* to −∞for 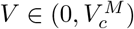, thus *f* (*V* ) has a unique zero point in 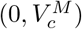, which we denote by *V*_0_. And *V* ∈ (0, *V*_0_).

Now let 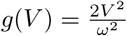 and *h*(*V* ) = *g*(*V* )*f* (*V* ), we get 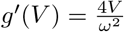, and

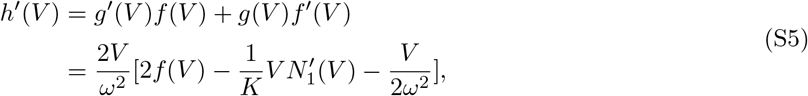

It’s easy to see that 2*f* (*V* ) monotonically decreases with *V* and 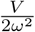 monotonically increases with *V* . To check the term 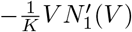, we compute 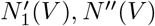 and get

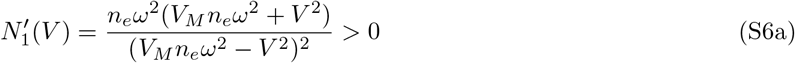

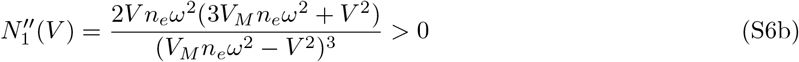

This means *N* ^*′*^(*V* ) is positive and monotonically increases with *V*, thus the term 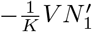 (*V* monotonically decrease with *V* . This means the function 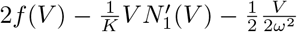 decreases monotonically from 2r > 0 to 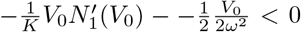, for *V* ∈ (0, *V*_0_ ), this means *h*(*V* ) first increases and then decreases, with a maximum reached at a point *V*_*c*_.

Thus, in order for a positive solution to exist, we must have *v*^2^ ≤ *h*(*V*_*c*_). This gives a bound for rate of moving optimum, which is consistent with classical theory.

### S2 Stability analysis

The question is then whether the solutions are then stable. To determine the local stability of the equilibrium (Δ^∗^, *V* ^∗^, *N* ^∗^), we compute the Jacobian matrix **J**:

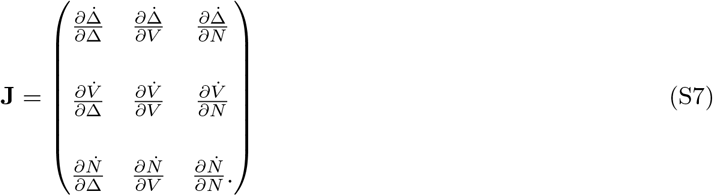

Evaluating the partial derivatives at the equilibrium:

- 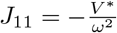
- 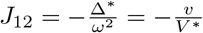
- 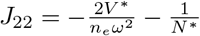
- 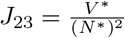
- 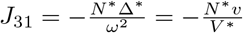
- 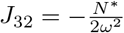
- 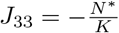

All other terms (*J*_13_, *J*_21_, *J*_33_) are zero. The resulting matrix is:

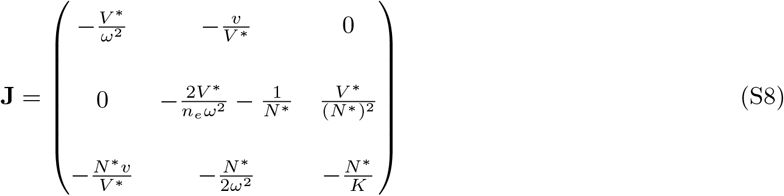

For the third-order system, the characteristic equation is given by *λ*^3^ + *a*_1_*λ*^2^ + *a*_2_*λ* + *a*_3_ = 0. Local asymptotic stability (Routh-Hurwitz stability criterion) requires:

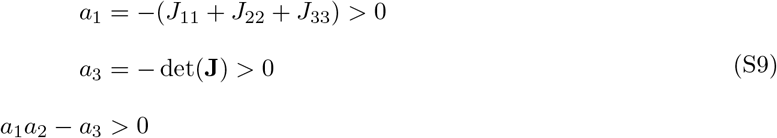

Notice that at the equilibrium *V* ^∗^, the first condition is always satisfied since *J*_11_, *J*_22_, *J*_33_ < 0. To check the second condition, we compute and simplify det(**J**), expanding det(**J**) along the third column

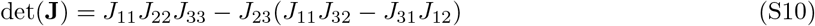

Substituting the components and we get

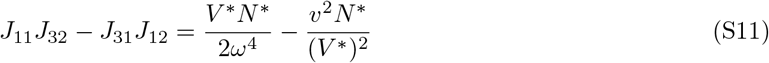

Thus

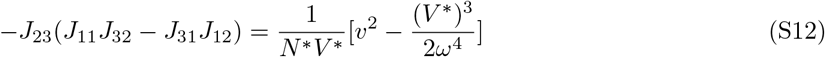

On the other hand, we have

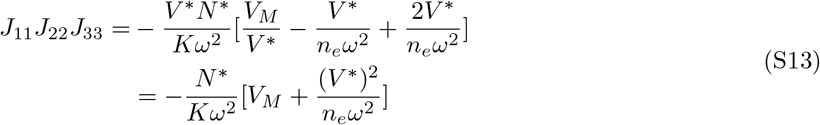

Take derivative on *h*(*V* ) at *V* ^∗^ and use the expression of *N* ^∗^ we get

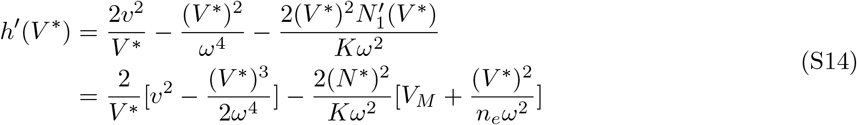

Comparing det(**J**) with 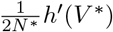 we get

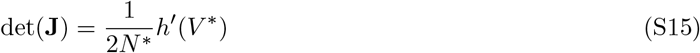

Now, when *v*^2^ < *h*(*V*_*c*_), *v*^2^ = *h*(*V* ) has two solutions, which we denote by *V*_1_ < *V*_*c*_ < *V*_2_. Since *h*(*V* ) first increases and then decreases, we know *h*^*′*^(*V*_1_) > 0 and *h*^*′*^(*V*_2_) < 0, and thus det(**J**) is positive at *V*_1_ and negative at *V*_2_, thus the only possible positive stable equilibrium is *V* = *V*_2_.

Now we go to the third condition for stability analysis, i.e. *a*_1_*a*_2_ − *a*_3_ > 0. To do this, we first write out *a*_1_, *a*_2_ and *a*_3_,

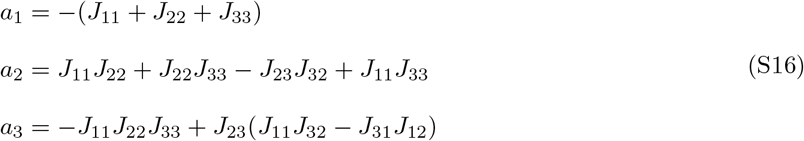

We then write *a*_1_*a*_2_ − *a*_3_ as two parts, *P*_1_ and *P*_2_, where *P*_1_ can be written as following

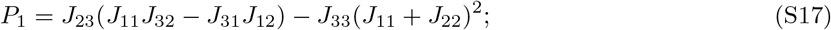

and *P*_2_ can be written as following

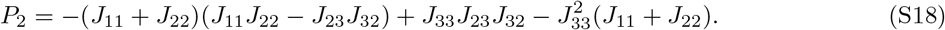

We first notice that *P*_2_ is always positive. This can be easily proved using the fact that *J*_11_ < 0, *J*_22_ < 0, *J*_33_ < 0, *J*_23_ > 0, *J*_32_ < 0. Thus we only need to check *P*_1_, for *P*_1_, we have

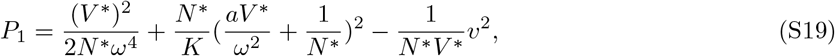

where 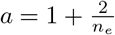

From *h*^*′*^(*V* ^∗^) < 0 at *V* ^∗^ = *V*_2_, we know

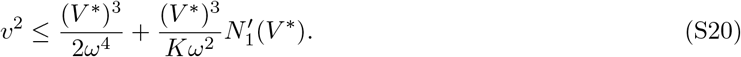

This means

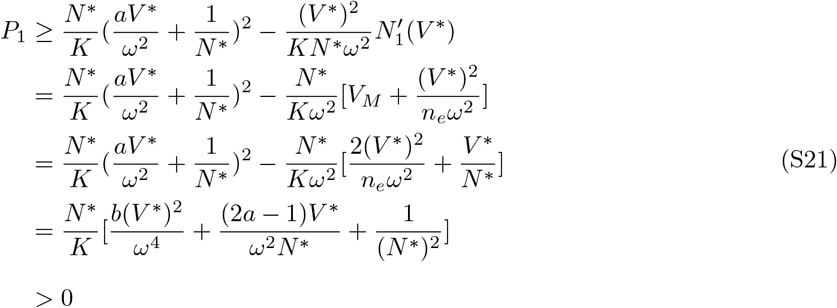

where 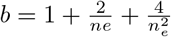

We thus have proven that the positive equilibrium *V* ^∗^ = *V*_2_ is always locally stable as long as it exists. As a brief summary, we have proven that when *v*^2^ *> h*(*V*_*c*_), the population will always go extinct. When *v*^2^ ≤ *h*(*V*_*c*_), both extinction and the positive equilibrium will be stable, and we have bistability in this case.

## References

Agashe, D., Falk, J. J., & Bolnick, D. I. (2011). Effects of founding genetic variation on adaptation to a novel resource. Evolution, 65 (9), 2481–2491.

Alexander, H. K., Martin, G., Martin, O. Y., & Bonhoeffer, S. (2014). Evolutionary rescue: Linking theory for conservation and medicine. Evolutionary Applications, 7 (10), 1161–1179.

Anstett, D. N., Anstett, J., Sheth, S. N., Moxley, D. R., Branch, H. A., Jahani, M., Angert, A. L., et al. (2026). Rapid evolution predicts demographic recovery after extreme drought. Science, 391 (6790), 1172–1176.

Bell, G. (2017). Evolutionary rescue. Annual Review of Ecology, Evolution, and Systematics, 48, 605–627.

Bell, G., & Gonzalez, A. (2011). Adaptation and evolutionary rescue in metapopulations experiencing environmental deterioration. Science, 332 (6035), 1327–1330.

Bell, G., & Gonzalez, A. (2009). Evolutionary rescue can prevent extinction following environmental change. Ecology Letters, 12 (9), 942–948.

Bulmer, M. G. (1971). The effect of selection on genetic variability. The American Naturalist, 105 (943), 201–211.

Bürger, R., & Lynch, M. (1995). Evolution and extinction in a changing environment: A quantitative-genetic analysis. Evolution, 49 (1), 151–163.

Gienapp, P., Lof, M., Reed, T. E., McNamara, J., Verhulst, S., & Visser, M. E. (2013). Predicting demographically sustainable rates of adaptation: Can great tit breeding time keep pace with climate change? Philosophical Transactions of the Royal Society B: Biological Sciences, 368 (1610), 20120289.

Gomulkiewicz, R., & Houle, D. (2009). Demographic and genetic constraints on evolution. The American Naturalist, 174 (6), E218–sE229.

Gonzalez, A., & Bell, G. (2013). Evolutionary rescue and adaptation to abrupt environmental change depends upon the history of stress. Philosophical Transactions of the Royal Society B: Biological Sciences, 368 (1610), 20120079.

Gonzalez, A., Ronce, O., Ferriere, R., & Hochberg, M. E. (2013). Evolutionary rescue: An emerging focus at the intersection between ecology and evolution. Philosophical Transactions of the Royal Society B: Biological Sciences, 368 (1610), 20120404.

Hao, Y. Q., Brockhurst, M. A., Petchey, O. L., & Zhang, Q. G. (2015). Evolutionary rescue can be impeded by temporary environmental amelioration. Ecology letters, 18 (9), 892–898.

Hoffmann, A. A., & Sgró, C. M. (2011). Climate change and evolutionary adaptation. Nature, 470 (7335), 479–485.

Höllinger, I., Pennings, P. S., & Hermisson, J. (2019). Polygenic adaptation: From sweeps to subtle frequency shifts. PLoS Genetics, 15 (3), e1008035.

Kimura, M. (1965a). Attainment of quasi linkage equilibrium when gene frequencies are changing by natural selection. Genetics, 52 (5), 875–890.

Kimura, M. (1965b). A stochastic model concerning the maintenance of genetic variability in quantitative characters. Proceedings of the National Academy of Sciences, 54 (3), 731–736.

Klausmeier, C. A., Osmond, M. M., Kremer, C. T., & Litchman, E. (2020). Ecological limits to evolutionary rescue. Philosophical Transactions of the Royal Society B: Biological Sciences, 375 (1814), 20190453.

Kopp, M., & Matuszewski, S. (2014). Rapid evolution of quantitative traits: Theoretical perspectives. Evolutionary Applications, 7 (1), 169–191.

Kopp, M., & Hermisson, J. (2009). The genetic basis of phenotypic adaptation i: Fixation of beneficial mutations in the moving optimum model. Genetics, 182 (1), 233–249.

Lachapelle, J., & Bell, G. (2012). Evolutionary rescue of sexual and asexual populations in a deteriorating environment. Evolution, 66 (11), 3508–3518.

Lachapelle, J., Colegrave, N., & Bell, G. (2017). The effect of selection history on extinction risk during severe environmental change. Journal of Evolutionary Biology, 30 (10), 1872–1883.

Lande, R. (1979). Quantitative genetic analysis of multivariate evolution, applied to brain: Body size allometry. Evolution, 33 (1), 402–416.

Lande, R. (1993). Risks of population extinction from demographic and environmental stochasticity and random catastrophes. The American Naturalist, 142 (6), 911–927.

Lande, R., & Shannon, S. (1996). The role of genetic variation in adaptation and population persistence in a changing environment. Evolution, 50 (1), 434–437.

Lande, R. (1975). The maintenance of genetic variability by mutation in a polygenic character with linked loci. Genetics Research, 26 (3), 221–235.

Lindsey, H. A., Gallie, J., Taylor, S., & Kerr, B. (2013). Evolutionary rescue from extinction is contingent on a lower rate of environmental change. Nature, 494 (7438), 463–467.

Lynch, M., & Lande, R. (1993). Evolution and extinction in response to environmental change. In Biotic interactions and global change (pp. 234–250, Vol. 49). Sinauer Associates Inc.

Matuszewski, S., Hermisson, J., & Kopp, M. (2015). Catch me if you can: Adaptation from standing genetic variation to a moving phenotypic optimum. Genetics, 200 (4), 1255–1274.

Negm, S., & Veller, C. (2026). The effect of long-range linkage disequilibrium on allele-frequency dynamics under stabilizing selection. PLOS Genetics, 22 (3), e1012035.

Nordstrom, S. W., Hufbauer, R. A., Olazcuaga, L., Durkee, L. F., & Melbourne, B. A. (2023). How density dependence, genetic erosion and the extinction vortex impact evolutionary rescue. Proceedings of the Royal Society B: Biological Sciences, 290 (2011), 20231228.

Osmond, M. M., & Klausmeier, C. A. (2017). An evolutionary tipping point in a changing environment. Evolution, 71 (12), 2930–2941.

Osmond, M. M., Otto, S. P., & Martin, G. (2020). Genetic paths to evolutionary rescue and the distribution of fitness effects along them. Genetics, 214 (2), 493–510.

Pahujani, S., Stetter, M. G., & Krug, J. (2026). Adaptive dynamics of quantitative traits in a steadily changing environment. Genetics, iyag168.

Parmesan, C. (2006). Ecological and evolutionary responses to recent climate change. Annual Review of Ecology, Evolution, and Systematics, 37 (1), 637–669.

Radchuk, V., Reed, T., Teplitsky, C., van de Pol, M., Charmantier, A., Hassall, C., Adamík, P., Adriaensen, F., Ahola, M. P., Arcese, P., et al. (2019). Adaptive responses of animals to climate change are most likely insufficient. Nature Communications, 10 (1), 3109.

Ramsayer, J., Kaltz, O., & Hochberg, M. E. (2013). Evolutionary rescue in populations of Pseudomonas fluorescens across an antibiotic gradient. Evolutionary Applications, 6 (4), 608–616.

Samani, P., & Bell, G. (2010). Adaptation of experimental yeast populations to stressful conditions in relation to population size. Journal of Evolutionary Biology, 23 (4), 791–796.

Samani, P., & Bell, G. (2016). The ghosts of selection past reduces the probability of plastic rescue but increases the likelihood of evolutionary rescue to novel stressors in experimental populations of wild yeast. Ecology Letters, 19 (3), 289–298.

Servedio, M. R., Brandvain, Y., Dhole, S., Fitzpatrick, C. L., Goldberg, E. E., Stern, C. A., Van Cleve, J., & Yeh, D. J. (2014). Not just a theory—the utility of mathematical models in evolutionary biology. PLoS biology, 12 (12), e1002017.

Turelli, M., & Barton, N. H. (1990). Dynamics of polygenic characters under selection. Theoretical Population Biology, 38 (1), 1–57.

Uecker, H., Osmond, M. M., Alejandre, C., Berríos-Caro, E., Clement, D. T., Czuppon, P., Dekens, L., Deraje, P., Draghi, J. A., Hasan, A., Holt, R. D., Kuosmanen, T., Longcamp, A., Marrec, L., Martin, G., Nyhoegen, C., Orive, M. E., Ravi Kumar, V., Ronce, O., … Yamamichi, M. (2026). Modeling evolutionary rescue. EcoEvoRxiv. 10.32942/X25D4T

Urban, M. C. (2015). Accelerating extinction risk from climate change. Science, 348 (6234), 571–573.

Willi, Y., & Hoffmann, A. A. (2009). Demographic factors and genetic variation influence population persistence under environmental change. Journal of Evolutionary Biology, 22 (1), 124–133.

Xu, K., & Osmond, M. M. (2026). When does the probability of evolutionary rescue increase with the strength of selection despite a potential demographic cost? The American Naturalist, 207 (5), 615–626.

Xu, K., Vision, T. J., & Servedio, M. R. (2023). Evolutionary rescue under demographic and environmental stochasticity. Journal of Evolutionary Biology, 36 (10), 1525–1538.

Zhou, D. H., & Zhang, Q. G. (2024). Loss and recovery of ecological diversity associated with evolutionary rescue in abruptly and gradually deteriorating environments. Evolution, 78 (4), 768–777.

